# Cancer cell exosomes can initiate malignant cell transformation

**DOI:** 10.1101/360982

**Authors:** Karoliina Stefanius, Kelly A. Servage, Marcela de Souza Santos, Jason Toombs, Hillery Fields Gray, Suneeta Chimalapati, Min S. Kim, Rolf A. Brekken, Kim Orth

## Abstract

Cancer evolves through a multistep process that occurs by the temporal accumulation of genetic mutations mediated by intracellular and extracellular cues. We observe that exosomes isolated from pancreatic cancer cells, but not normal pancreatic cells, can initiate the first step of malignant cell transformation. Injection of exosome-initiated transformed cells into mice results in aggressive tumor growth. Using proteomic profiling and DNA sequencing of exosome-treated and transformed cells, we show that cancer cell exosomes act as a classic initiator by causing random genetic changes in recipient cells. Our studies provide new insight into a function of cancer cell exosomes and how they might specifically contribute to orchestrated local cell transformation.

**One Sentence Summary:** Exosomes function as an *initiator* of tumor formation.

## Main Text

Within the tumor microenvironment, a dynamic molecular communication between tumor and surrounding stromal cells is a well-recognized feature of cancer progression (*1, 2*). The conversation at the primary tumor site, as well as at distant locations, is mediated through many secreted factors including exosomes; small (30-150nm) secreted extracellular vesicles (sEV) shed by normal and malignant cells (*3-6*). Interest in exosomes and their mechanism(s) of biogenesis and function has emerged as a promising, yet controversial, field of research. Based on multi-omic studies, these sEV are known to carry heterogeneous cargo composed of proteins, metabolites, genetic material (DNA and microRNAs), and lipids (*7-12*). They are selectively packaged and transferred into recipient cells acting as vehicles in intercellular communication in normal physiological and pathological conditions (*13*). A growing body of evidence shows sEV are crucial in shaping the local tumor microenvironment to promote cancer progression by advancing tumor metastasis (*4, 14*). Although there is a major emphasis on describing the function of exosomes in metastasis and interactions between local tumor microenvironments, less effort has been invested in analyzing their possible contribution to transforming a normal cell into a malignant cell. Since pancreatic cancer is a lethal metastatic disease that lacks efficient curative treatment, we raised the question of whether sEV, secreted by pancreatic cancer cells, participate directly in the process of cellular transformation (*15*).

Malignant transformation of a normal cell occurs in a stepwise fashion. Initial changes induced by environmental exposure, even with relatively low doses or genetic cues, can result in the reprogramming of a normal cell to a less differentiated state that is receptive to additional genetic alterations resulting in uncontrolled growth and ultimately cancer. The classic 2-stage *in vitro* cell transformation assay is a tiered system for transformation that was created for screening potential carcinogenic factors (*16-18*). In this system, cells are first treated with a suspected carcinogen, called an *initiator*, which causes random genetic changes in a pool of normal cells. Subsequently, these *initiated* cells are exposed to a *promoter*, which induces further genetic alterations resulting in cell transformation and observed as foci on a cell culture plate (**Fig. 1A**) (*18, 19*). This 2-stage assay provides sensitivity in detecting a wider range of *initiating* agents that may not show obvious transforming activity without a *promote*r (*18*). With the recent indications that exosomes are emerging contributors to normal physiology and tumor promotion, we asked a simple question: can exosomes function as an *initiator* or *promoter* during the transformation of a normal cell to a cancer cell?

**Figure 1.**
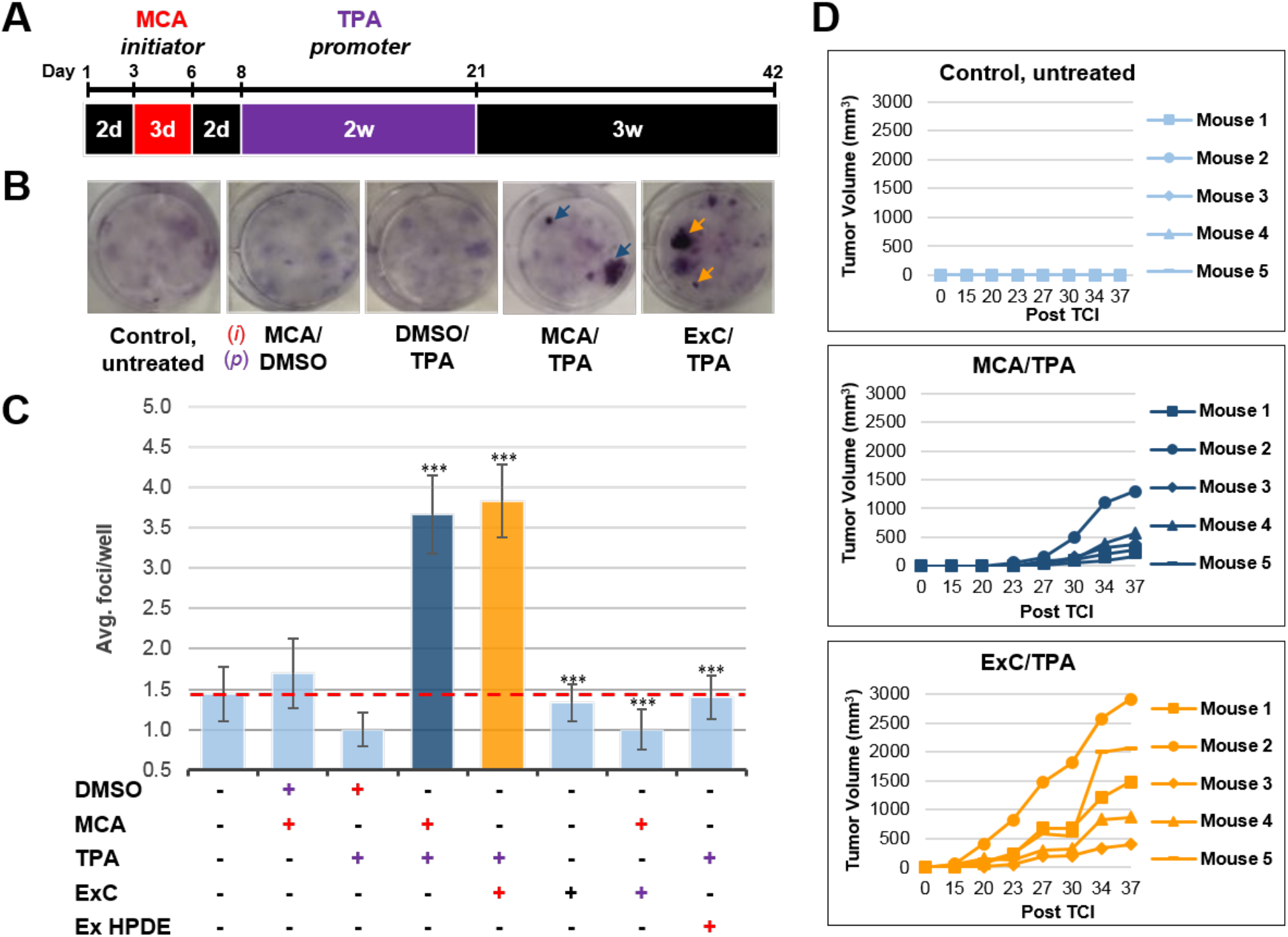
Pancreatic cancer cell exosomes function as an *initiator* in malignant cell transformation. **(A)** In the classic 2-stage cell transformation assay used, NIH/3T3 cells are treated with the tumor *initiator* MCA for three days (Days 3-6) and the tumor *promoter* TPA for two weeks (Days 8-21). After 42 days, cells are fixed with methanol and stained with Crystal Violet for malignant foci counting. **(B)** Representative images of stained cells showing foci formation (arrows) from untreated cells and cells treated with MCA/DMSO, DMSO/TPA, MCA/TPA, or Capan-2 exosomes (ExC)/TPA (*initiator-promoter*). **(C)** Quantification of foci formed after cell transformation. The average foci/well were determined via double-blind counting as described in methods. The red dashed line represents the established level of background foci present in untreated cells. Only MCA/TPA (p=0.008) and ExC/TPA (p=0.0002) treatments resulted in increased foci formation above background. Asterisks indicate significant differences from either control treatment or MCA/TPA treatments as determined by unpaired, two-tailed t-test with Welch’s correction (*p<0.05; **p<0.01; ***p<0.001; ****p<0.0001). Red (+) = *initiator*; purple (+) = *promoter;* HPDE exosomes (Ex HPDE) **(D)** Subcutaneous injections of NIH/3T3 cells, MCA/TPA transformed cell foci, or ExC/TPA transformed cell foci into mice (n=5) at a concentration of 2.5−10^6^ cells. Tumor growth was tracked by measuring tumor volume 2x/week for 37 days post injection (Post TCI).

To address this question, we utilized the classic cell transformation assay with NIH/3T3 cells using 3-methylcholanthrene (MCA) and tetradecanoyl phorbol acetate (TPA) as controls for an *initiator* and *promoter*, respectively (**Fig. 1A**) (*18, 20*). Untreated cells or cells treated with MCA or TPA alone showed low levels of background foci, as defined by criteria on focus scoring (**Fig. 1B and C, table S2**) (*19*). By contrast, cells treated for three days with MCA (*initiator*) followed by two weeks of TPA treatment (*promoter*) resulted in the consistent formation of 3-4 foci per well (**Fig. 1B and C, table S2**). Therefore, only after treatment with both an *initiator* and *promoter* do we observe an increased number of foci above background levels (**Fig. 1B and C**).

To investigate whether exosomes can act as an *initiator* or *promoter*, we isolated exosomes from a pancreatic cancer cell line, Capan-2, using an established ultrafiltration protocol (*21*). Prior to treating cells with exosomes, we validated the purification by following a published minimal criteria protocol, with follow-up assays including electron microscopy, mass spectrometry, and immunoblot analysis on biological triplicates, to confirm rigor and reproducibility of our exosome preparations (**fig. S1A to G, table S1**) (*22, 23*). Treatment of cells with only Capan-2 exosomes for the duration of the *initiation* and *promotion* steps (three weeks) showed a low level of background transformation activity (**Fig. 1C, table S2**). Similarly, when testing Capan-2 exosomes as a *promoter*, with the *initiator* MCA, we also observed background transformation activity comparable to control treatments (**Fig. 1C, table S2**). However, when Capan-2 exosomes were used as the *initiator*, with the *promoter* TPA, we observed comparable foci formation (3-4 foci per well) to the traditional MCA/TPA treatment (**Fig. 1C, table S2**). Therefore, Capan-2 exosomes can function as an *initiator*, when combined with the *promoter* TPA, to induce cellular transformation.

We repeated these experiments with exosomes isolated from other pancreatic cancer cells, including Mia PACA-2, Panc-1, and BxPC-3, and exosomes isolated from normal immortalized pancreatic cells (HPDE) (**Fig. 1C, fig. S2A, table S3**). We observed that, similar to Capan-2 exosomes, Mia PACA-2 and Panc-1 cancer cell exosomes can function as an *initiator*, but not alone or as a *promoter*, to induce cellular transformation (**Fig. 1C, fig. S2A**). These three pancreatic cell lines (Capan-2, Mia PACA-2, and Panc-1) contain oncogenic mutations in the Ras gene, whereas, the BxPC-3 cell line contains a wild type Ras gene. Considering exosomes with oncogenic *KRAS* can act as an *initiator*, we tested whether BxPC-3 has the same *initiator* activity (**fig. S2A**). We conclude that the observed exosome-mediated *initiator* activity is independent of the state of the Ras gene, as all four pancreatic cancer cell lines, including BxPC-3, have the ability to function as an *initiator* (**fig. S2A**). Importantly, exosomes isolated from the normal immortalized pancreatic cell line HPDE were unable to induce cellular transformation when used as an *initiator* in the assay (**Fig. 1C**). Therefore, pancreatic cancer cell exosomes, but not exosomes isolated from normal pancreatic cells, can *initiate* cellular transformation.

A common characteristic of *initiator*s in the cellular transformation assay is that the *initiator* can function with multiple *promoters* to induce cell transformation (*24, 25*). To test whether the pancreatic cancer cell exosomes could act as an *initiator* with other *promoters*, we replaced TPA with another common *promoter*, cadmium chloride (CdCl_2_), in the transformation assay (*24, 26, 27*). We observed that cells treated with either MCA or Capan-2 exosomes as an *initiator* followed by treatment with CdCl_2_ as a *promoter* resulted in transformation of cells, similar to what was observed when using TPA as the *promoter* (**fig. S2B, table S4**). Treatment of cells with CdCl_2_ alone resulted in background levels of foci, reiterating the fact that cell transformation is dependent on both *initiation* and *promotion*. These results indicate that exosomes isolated from pancreatic cancer cells are indeed acting as general *initiators* in the transformation assay and are not dependent on a specific *promoter*.

An important step of assessing the tumorigenic property of transformed cells is their ability to form tumors *in vivo*. To determine whether exosome-mediated transformed cells have the capacity to form tumors when injected subcutaneously into immunocompromised mice, we isolated and expanded foci cells from the MCA/TPA and Capan-2 exosome/TPA experiments. The cells from these foci were then injected into NSG (NOD scid gamma) mice at three different concentrations and the mice were followed for the presence of measurable tumor formation (**Fig 1D, fig. S3A**). As a control, non-transformed NIH/3T3 cells were injected into mice at the highest concentration. Tumor growth was observed in mice injected with cells from MCA/TPA and Capan-2 exosome/TPA foci, but not with NIH/3T3 cells (**Fig. 1D**). The appearance of tumors correlated with the number of cells injected into mice such that higher concentration of cells resulted in faster growth and ultimately larger tumors (**Fig. 1D, fig. S3A**). Histological analysis confirmed that the tumors are fibrosarcomas, as expected from transformed cells of mesenchymal origin (**fig. S3B**). To support the results obtained from *in vitro* experiments with other cancer cell exosomes, we performed the same *in vivo* studies in mice using Mia PACA-2 exosome/TPA, Panc-1 exosome/TPA, and BxPC-3 exosome/TPA foci. We observed that cells from these cancer cell exosome-*initiated* foci also formed tumors when injected into mice (**fig. S2C**). These studies clearly demonstrate that pancreatic cancer cell exosomes, independent of the state of the Ras gene, can function as an *initiator* to produce transformed cells that can induce tumor growth when injected into mice.

We note here that the exosomes used in our assays are derived from human cancer cell lines whereas the “normal” cells used in the cell transformation assay are NIH/3T3 cells from murine origin. Previous studies have shown highly conserved molecular functions across human and mouse, including the regulation of cell division, DNA replication, and DNA repair (*28*). Moreover, since some of the microRNAs are conserved between human and mice (*29*) and show a proclivity to change signaling in a treated cell, we first analyzed microRNA associated with cancer cell (Capan-2) and normal cell (HPDE) exosomes for differences. Analysis showed that a large majority of microRNA present were found to be upregulated in cancer cell exosomes compared to normal cell exosomes (**fig. S4, table S5**). Considering this, we analyzed whether the three-day *initiator* treatment with the human cancer cell exosomes could globally change signaling in the murine NIH/3T3 cells used in the cell transformation assay.

To asses this, NIH/3T3 cells were either left untreated or treated for three days with Capan-2 exosomes or MCA as the *initiator* followed by two days of recovery. Cells were harvested, and total protein was analyzed by mass spectrometry. As expected, NIH/3T3 cells left untreated or treated with MCA showed no marked global changes in protein content (**Fig. 2A**). MCA functions by incorporating random mutations to *initiate* cellular transformation (*30*) and, therefore, it is expected that changes to proteins in any one cell among the total population would not be feasible to detect. Interestingly, we observed a similar proteomic profile of NIH/3T3 cells treated for three days with Capan-2 exosomes, showing that initiation alone is not altering the protein composition of cells (**Fig. 2A**). In retrospect, these observations are not so surprising because it is known that exosomes do not uniformly contain the same pool of proteins, lipids, metabolites, or microRNAs, but rather each exosome contains a unique repertoire of biological molecules (*8, 11, 12*). Therefore, as observed with other *initiators* such as MCA, cancer cell exosomes are likely causing random molecular changes across the population of treated NIH/3T3 cells that cannot be detected amongst the overall protein composition of cells.

**Figure 2.**
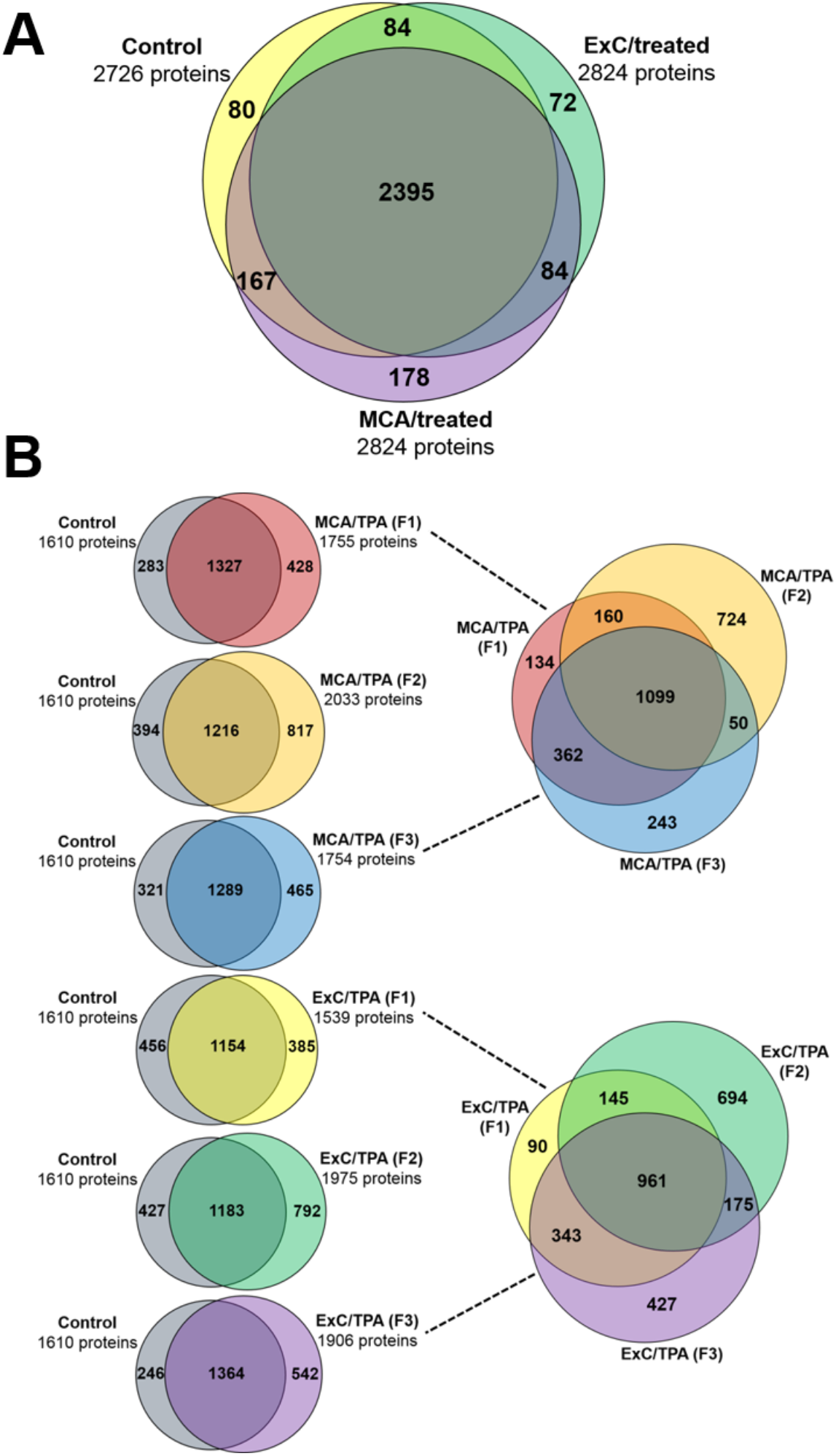
Proteomic profiling of initiated and transformed NIH/3T3 cells via mass spectrometry analysis. **(A)** Comparison of proteins found in cells left untreated (control) versus a 3-day initiation treatment with either MCA or Capan-2 (ExC) exosomes. For each condition, the results from three biological replicates were combined. **(B)** Comparison of proteins found in transformed cells resulting from treatment with both an *initiator* and *promoter*. Three separate foci (F1, F2, F3) of MCA/TPA transformed cells and ExC/TPA transformed cells were compared. Results from three biological replicates were combined for each of the six foci, as well as the untreated control.

Based on these observations, we propose that cancer cell exosomes have the capacity to act as an *initiator* by incorporating random changes to mediate the first step in cellular transformation. We predict that detection of these single changes within the total population of MCA- or cancer cell exosome-treated cells would be like trying to find a “needle in a haystack”. Therefore, we turned to the foci that are formed during the 2-stage cellular transformation assay to analyze and compare proteomic and exome profiles of untreated NIH/3T3 cells, three MCA/TPA foci, and three Capan-2 exosome/TPA foci. Analyses of the proteomic profiles of these samples revealed that the transformed cells had diverged significantly from the control NIH/3T3 cells (**Fig. 2B**). Venn diagrams comparing the three MCA/TPA foci and the three Capan-2 exosome/TPA foci revealed unique differences in the protein composition of each of the transformed cell samples, indicating that the *initiator* and *promoter* events combined to result in diverse changes to proteins in foci of transformed cells. Further, it appears that these changes are not uniform when comparing foci formed from the same treatment (**Fig. 2B**).

Exome-seq analysis was performed in triplicates for MCA/TPA and Capan-2 exosome/TPA foci to better understand the genetic mechanism of cellular transformation. No common mutations were found among 190 known oncogenes (**Data S5.**) supporting the proposal that random events accumulated to induce a transformed phenotype. Clustering analysis (**Fig. 3A**) (*31*) of the samples also indicates no clear relationship between conditions (**Fig. 3B**). When using MutaGene (*32*) to investigate the specific type of nucleotide changes in the six foci compared to NIH/3T3 untreated cells, the mutational signature Cosmic 20 (**Fig. 3C**), associated with defective mismatch repair and microsatellite instability, was observed across all samples. Interestingly, when mismatch repair genes were analyzed in these foci samples, missense and nonsense mutations were found to be encoded by these transformed cells, indicating the genetic source for the mutation profile observed (**Table 1**). Furthermore, none of the foci contain the same mutations. Since all six foci where derived from treatment with a common *promoter*, it is tempting to propose that future studies might reveal that a *promoter* drives the selection for the mutator phenotype.

**Figure 3.**
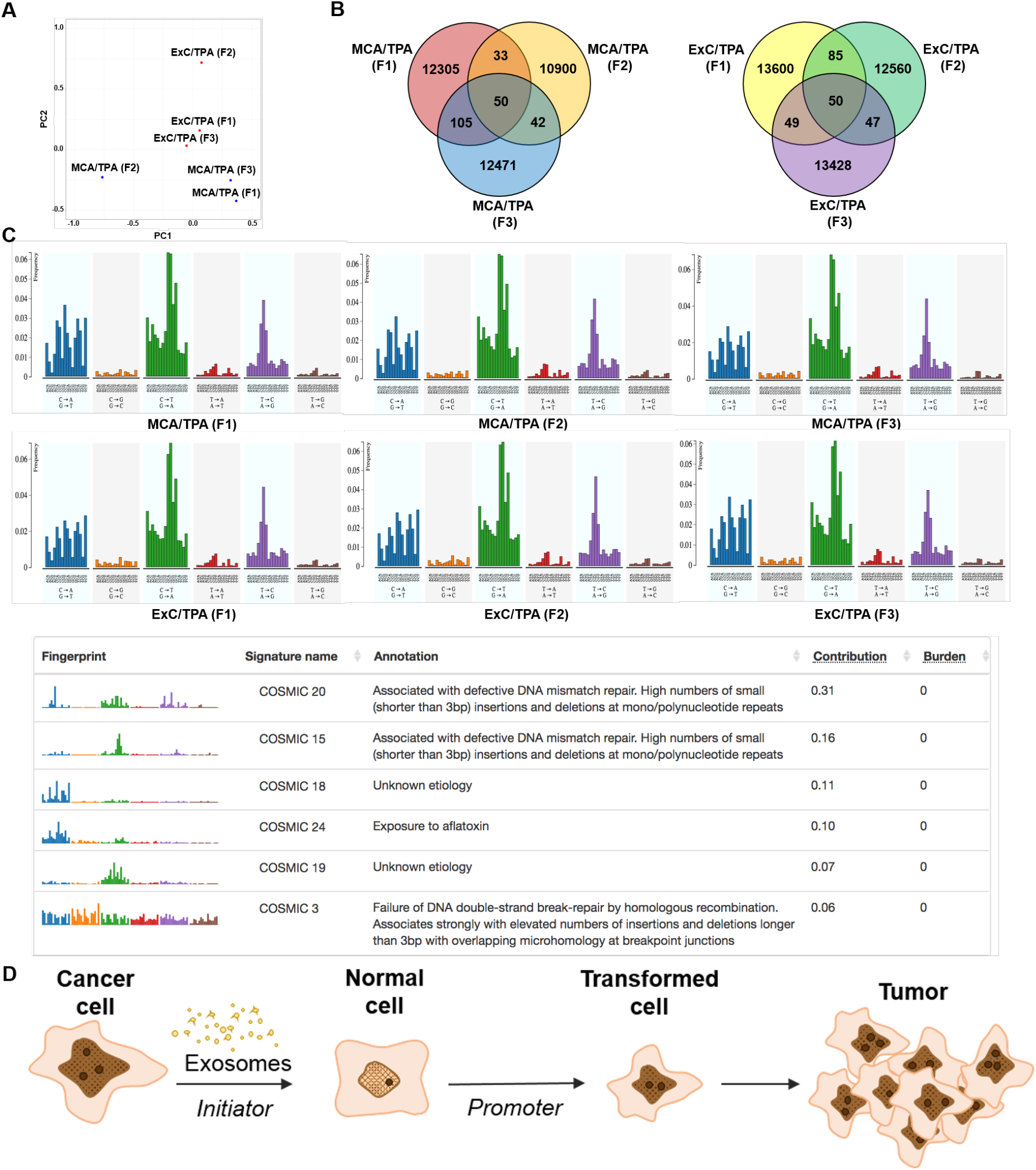
DNA Sequencing of transformed NIH/3T3 cells. **(A)** Principle component analysis showing relationship of MCA/TPA and ExC/TPA transformed cell samples based on comparison of exome-seq variant data using PLINK’s identity-by-state (IBS) estimates. **(B)** Total number of variants found in MCA/TPA and ExC/TPA samples using exome-seq. **(C)** Mutational profiles of six samples (MCA/TPA F1, F2, F3 and ExC/TPA F1, F2, F3) sent for DNA sequencing along with identity of most similar profile (COSMIC 20) as annotated by MutaGene. **(D)** Schematic model of exosome mediated transformation. Exosomes secreted by cancer cells are taken up by normal cells and have the capacity to act as an *initiator* by incorporating random changes into the recipient cell genome. These initiated cells, when exposed to a *promoter*, can be induced by further alterations to a transformed state that has the ability to grow into a malignant tumor.

**Table 1.** Non-synonymous variants found in mismatch repair associated genes (Msh2, Msh3, Msh6, Pms1, Pms2, Mlh1) across all six samples.

| Gene | Chr | HGVS.c | MCA/TPA (F1) | MCA/TPA (F2) | MCA/TPA (F3) | ExC/TPA (F1) | ExC/TPA (F2) | ExC/TPA (F3) | Annotation |
| --- | --- | --- | --- | --- | --- | --- | --- | --- | --- |
| *Msh3* | 13 | c.2137G>T |  |  |  |  |  | p.Glu713* | Stop gained |
| *Msh2* | 17 | c.1054G>C |  |  | p.Asp352His |  |  |  | Missense variant |
| *Msh2* | 17 | c.1904A>C |  |  |  | p.Lys635Thr |  |  | Missense variant |
| *Msh2* | 17 | c.2086C>T |  |  |  | p.Pro696Ser |  |  | Missense variant |
| *Msh2* | 17 | c.2215G>A | p.Ala739Thr |  |  |  |  |  | Missense variant |
| *Msh6* | 17 | c.67G>A |  |  |  |  |  | p.Ala23Thr | Missense variant |
| *Msh6* | 17 | c.1703T>C |  |  |  |  |  | p.Phe568Ser | Missense variant |
| *Msh6* | 17 | c.2258G>A |  |  |  |  |  | p.Gly753Glu | Missense variant |
| *Pms2* | 5 | c.323G>A |  |  |  |  |  | p.Gly108Glu | Missense variant |
| *Pms2* | 5 | c.1169G>A<br>c.2363G>A |  |  |  |  |  | p.Ser390Asn<br>p.Ser788Asn | Missense variant |
| *Mlh1* | 9 | c.226G>T |  | p.Val76Leu |  |  |  |  | Missense variant |
| *Pms1* | 1 | c.2087A>G |  |  |  | p.Lys696Arg |  |  | Missense variant |
| *Pms1* | 1 | c.1921A>G |  |  |  |  | p.Arg641Gly |  | Missense variant |
| *Pms1* | 1 | c.1686G>T |  |  |  |  | p.Met562Ile |  | Missense variant |
| *Pms1* | 1 | c.1530G>C |  |  |  |  |  | p.Lys510Asn | Missense variant |
| *Pms1* | 1 | c.1469G>T |  | p.Gly490Val |  |  |  |  | Missense variant |

Herein we observe that exosomes from pancreatic cancer cell lines can *initiate* the transformation of normal cells (**Fig. 3D**). Recently, analyses of neoplastic and carcinoma cells from pancreatic cancer patients demonstrated that adjacent cells from the same patient contain unique driver mutations, supporting the proposal that the cells in a tumor can be derived from independent transformation events as opposed to exclusive clonal events (*33*). These studies in combination with the observations in this report support a unique view of cellular transformation and how it is driven by the dynamic conversation between normal and cancer cells. Future studies are needed to determine how universal the cancer cell exosome *initiator* activity is and what specific factors from exosomes are contributing to cellular transformation.

## Acknowledgments

We thank members of the Orth lab for their helpful discussions and advice. We thank Diego Castrillon for his expert advice on tissue pathology and the UTSW Electron microscopy Core Facility.

## Author contributions

Conceiving the project: KS, KO and RAB. Conducting experiments: KS, SC. Performing mass spectrometric analysis of protein samples: KAS. Assisting with the experiments: HFG, JT. Animal studies: KS, HFG, JT. Data analysis: KS, KAS, MSS, MSK, KO. MicroRNA analysis: KS, MSK. Exome sequencing: KS, MSK. Writing the manuscript: KS, KAS, MSK, KO. Revising the manuscript: KS, KAS, MSS, HFG, SC, MSK, RAB, KO. Funding acquisition: K.O and RAB.

## Data and materials availability

All data is available in the main text or the supplementary materials.

## Supplementary Materials

Materials and Methods

Figures S1-S4

Tables S1-S5

Data S1-S5

References (*34-39*)

